# A Study of Microbial Diversity in a Biofertilizer Consortium

**DOI:** 10.1101/2023.05.15.540786

**Authors:** Cristóbal Hernández-Álvarez, Mariana Peimbert, Pedro Rodríguez-Martin, Dora Trejo-Aguilar, Luis D. Alcaraz

## Abstract

Biofertilizers supply living microorganisms to help plants grow and maintain their health. In this study, we examine the microbiome composition of a commercial biofertilizer that has been proven to promote plant growth. Using ITS and 16S rRNA gene sequence analyses, we describe the microbial communities of the biofertilizer, with 182 fungal species and 964 bacterial genera identified. The biofertilizer contains a variety of microorganisms that had been reported to enhance nutrient uptake, phytohormone production, stress tolerance, and pathogen resistance in plants. Plant roots created a microenvironment that boosted bacterial diversity but filtered fungal communities. We propose using plant roots as bioreactors to sustain dynamic environments that promote the proliferation of microorganisms with biofertilizer potential. However, preserving the fungal-inoculated substrate is crucial to maintain fungal diversity in the root fraction. The study suggests that bacteria grow close to plant roots, while root-associated fungi may be a subset of the substrate fungi. These findings indicate that the composition of the biofertilizer may be influenced by the selection of microorganisms associated with plant roots, which could have implications for the effectiveness of the biofertilizer in promoting plant growth.

## Introduction

Food demand has become a crucial concern for humanity’s future due to population growth, resource limitations, and climatic change [1,2]. The world’s dietary requirements will increase by 62 to 98% by 2050 [1]. Biotechnology can benefit food and agricultural production by using innovations to enhance the production process for animals, plants, and microorganisms [1]. Biofertilizers are biotechnological products that contain microorganisms applied to soil, seeds, or plant surfaces to promote vegetable growth [3,4]. Optimizing chemical fertilization in crops and transitioning to biofertilizer development is of utmost importance due to the environmental concerns associated with the excessive use of chemical fertilizers in agriculture. Using chemical fertilizers has led to the accumulation of nitrates in soil and water, which has disrupted the nitrogen cycle and contributed to the emission of nitrogen oxides in the atmosphere [5]. Excessive agricultural fertilizers also result in water pollution and greenhouse gas emissions, which can lead to catastrophic events such as the sargassum blooms affecting large areas in the Caribbean [6]. Similarly, excessive using phosphorus fertilizers in agriculture contributes to water pollution, and as phosphorus is an irreplaceable nutrient, it is essential to improve its assimilation in crops [7]. Microbes are capable of phosphorus uptake and assimilation of alternative P sources like phosphonates, and forming mycorrhizal interactions helps to provide a P source to the plant [8,9]. Favoring microbial consortia, like the ones included in some biofertilizers, could also impact the carbon cycle, creating a delicate balance between microbial metabolic activity in the plant-microbe interface and biogeochemistry. Such an approach could develop more effective and sustainable biofertilizers [10].

Biofertilizers put microbes to work, particularly in nitrogen fixation and phosphorus solubilization, and help to reduce the need for chemical fertilizers, which could mitigate climate change and improve soil health. Additionally, microbial consortia in biofertilizers could also impact the carbon cycle, developing more effective and sustainable biofertilizers [11]. For example, experiments using arbuscular mycorrhizal fungi (AMF) as biofertilizers reduced the application of external fertilizers, mainly phosphorus [12]. In the last decades, using soil microorganisms as biofertilizers have been a great success due to their benefits in promoting plant growth, pathogen control, increased quality, and crop yield enhancement [13,14]. Several companies sell biofertilizers based on plant-growth-promoting rhizobacteria (PGPR), *Rhizobium,* and mycorrhizal fungi. However, most commercial preparations advertise their products as general bacterial and fungal compositions, not declaring a species-level identification [15]. Biofertilizer production traditionally centers on screening, characterizing, and formulating single isolates with the desired plant-growth-promoting traits [16]. Nonetheless, the evidence suggests that bio-inoculants increase their effectiveness when using communities rather than single species [17,18].

AMF cultivation is challenging as they thrive as symbionts and thus limit *in vitro* cultivation [19]. Trap culture isolates AMF; it uses plants as hosts (traps) growth in soil mixed with sterile sand, usually in pots and use planta as baits to attract and host microbes [20–23]. It is also a technique that helps the long-term propagation of AMF [24]. Using plant microbial traps increases microbiome diversity by planting plant hosts into diverse soil samples from the field, then selecting the plant-interacting microorganisms [25]. Since their plant-beneficial effect depends on their capacity to colonize roots [26–28], understanding root-microbe interactions is essential for developing new biofertilizers. Trap culture propagation resembles the serial passage across generations in fresh media in experimental evolution. In other contexts like bioengineering, this is named long-term continuous culture [29]. Briefly, experimental evolution involves known ancestral populations (*e.g.*, microorganisms) propagated over time under selective pressures in managed conditions (*e.g.*, nutrients, heat) and then searching for the genetic basis of the adaptive traits; thus, fitness increases in the selective conditions [30]. Then, the genetic basis of the adaptations is revealed mainly by high-throughput sequencing [30].

AMF-based biofertilizers are produced through steps, including trap cultures from isolates, selection of suitable growth conditions (*e.g.*, low phosphorus medium), testing, propagating, and scale-up production [19]. Some biofertilizers use AMF communities (consortia for engineers) rather than a single AMF species [19]. In this study, we aimed to explore the role of plant roots in selecting microorganism communities responsible for promoting plant growth. We identified the species and genus-level composition of arbuscular mycorrhizal fungi (AMF) trap-based biofertilizers previously designed for this purpose [19]. Additionally, we described this biofertilizer’s overall microbial diversity, including its accompanying bacterial communities that were not originally intended to be included. Our study highlights the importance of understanding the microbial communities associated with biofertilizers, particularly those with plant-growth promotion capabilities.

## Materials and methods

### Biofertilizer production

The protocol for creating the biofertilizer discussed in this study is detailed in a publication by Trejo-Aguilar and Banuelos (2020) and is available commercially. The process involves sampling soil from natural ecosystems and searching for AMF spores of multiple species. Briefly, the soil is mixed with sterilized sand, and plants are used as microbial traps. Spores are recovered from the roots and preselected to promote the fast-growing of model plants in sterile soil. For massive propagation and consortium development, quartz sand and pumice is inoculated with host roots and soil. It is recommended to rotate plant hosts every four months, *Brachiaria brizantha, Crotalaria juncea*, and *Canavalia ensiformis*. After two to three months, watering is stopped, and the roots are harvested, dried, sieved, ground, and subjected to quality control inspection before packaging for commercial use. The resulting mixture contains sporulating microbes resistant to desiccation, and the biofertilizer’s activity is tested in fast-growing plants and their controls (Fig 1). In this study, we collected fresh root and substrate samples, stored them at −80°C, and extracted metagenomic DNA for microbiome sequencing.

**Fig 1.**
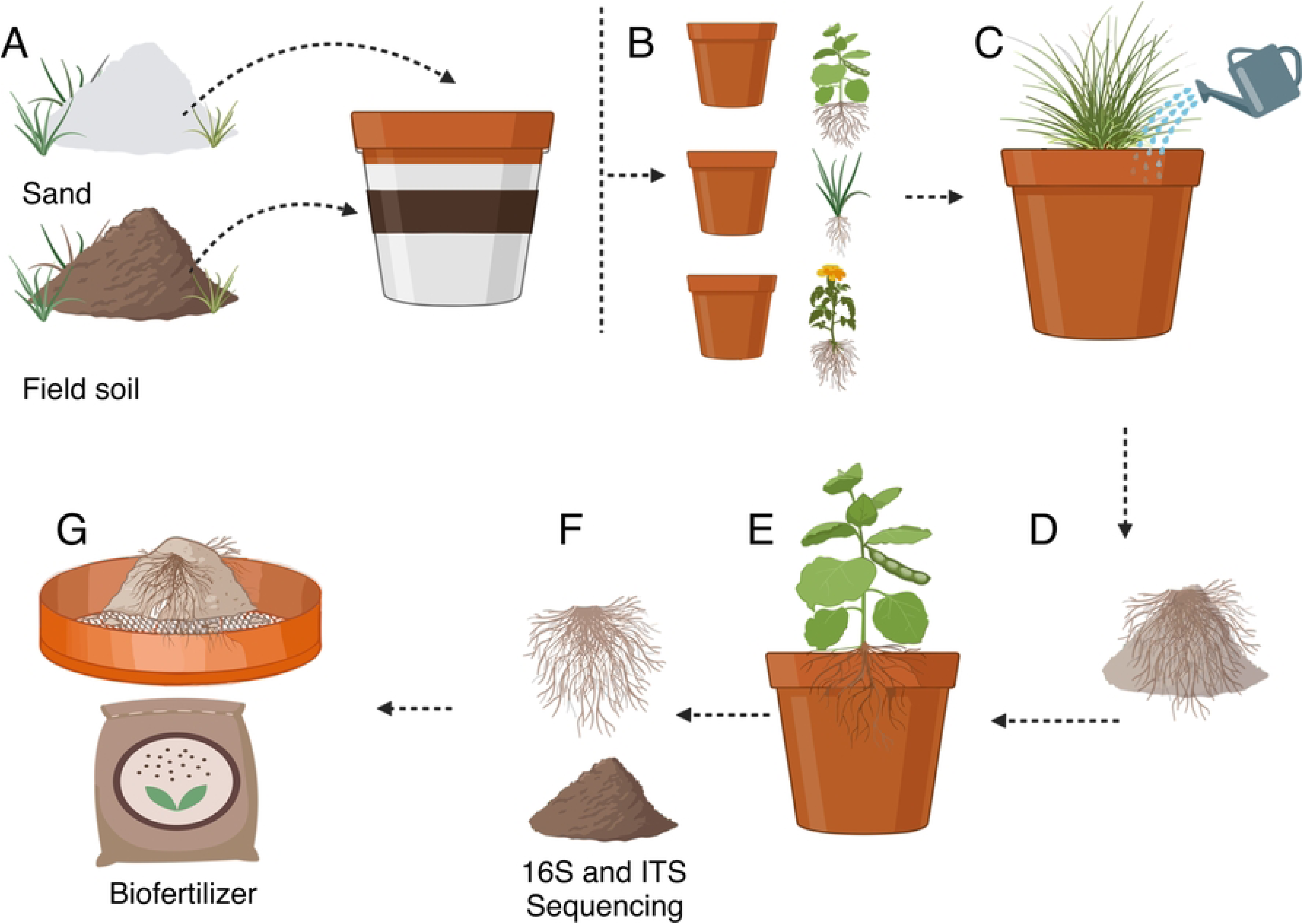
Biofertilizer production overview. (A) It starts with soil sampling from natural ecosystems, mixing the soil with sterilized sands, and (B) germinating plants to host arbuscular mycorrhizae and bacteria. (C) The resulting mixture of roots and substrate is then tested for its ability to promote plant growth. (D) Plant-growth promoters consortia are then selected and used as inoculum, roots and substrate, for (E) scale-up production and maintained through plant host rotation. (F) The substrate samples were collected for sequencing analysis before the drying process. (G) The roots and substrate are sun-dried and ground to produce the biofertilizer. The production process helps develop effective biofertilizers with diverse microbial communities that promote sustainable agriculture.

### Metagenomic DNA extraction and amplicon sequencing

We divided biofertilizer samples into four groups based on their source: fungi from the substrate, fungi from the roots, bacteria from the substrate, and bacteria from the roots. Following the manufacturer’s protocol, the metagenomic DNA was extracted using MoBio PowerSoil® DNA Isolation Kit (MoBio Laboratories, Solana Beach, CA, USA). Fungal diversity was assessed by PCR amplification of the ITS region using primers ITS1 (5’-TCGTCGGCAGCGTCAGATGTGTATAAGAGACAG-TCCGTAGGTGAACCTGCGG-3’) and ITS4 (5’-GTCTCGTGGGCTCGGAGATGTGTATAAGAGACAG-TCCTCCGCTTATTGATATGC-3’). For bacterial diversity, the 16S rRNA gene V3-V4 region was amplified by PCR using primers MiSeq341F (5’-TCGTCGGCAGCGTCA GATGTGTATAAGAGACAG-CCTACGGGNGGCWGCAG-3’) and MiSeq805R (5’-GTCTCGTGGGCTCGGAGATGTGTATAAGAGACAG-GACTACHVGGGTATCTAATCC-3’). Amplification of ITS region and 16S rRNA V3-V4 region followed the metagenomic DNA denaturation at 94°C for 3 min; then 20 denaturation cycles at 95°C for 30 s and extension at 72°C for 30 s. Next-generation sequencing of amplified regions was performed on Illumina® MiSeq TM (Illumina, San Diego, CA, USA) with 2 x 300 bp paired-end configuration at the *Laboratorio Nacional de Genómica para la Biodiversidad* (UGA-Langebio) for ITS region, and at the *Unidad de secuenciación Masiva* from *Biotechnology Institute*, *UNAM* for 16S rRNA gene.

### Sequence processing and data analyses

According to the previously reported protocol, we conducted ITS and 16S rRNA gene analyses (https://github.com/genomica-fciencias-unam/SOP; [31]). V3-V4 raw sequences were quality checked and trimmed 250 bp length using FASTX-Toolkit [32]. Sequence assembly was performed with Pandaseq [33] using a quality threshold of 0.95, a minimum length of 250 bp, and a minimum overlap of 15 bp. ITS1-ITS4 amplified sequences were pair-merged using CASPER [34]. Operational taxonomic units (OTUs) were clustered using cd-hit-est at 97% of identity [35]. Representative OTUs were extracted and taxonomically assigned using QIIME scripts [36]. ITS sequences were aligned against the UNITE v8.3 database [37] and 16S rRNA sequences against the SILVA v128 database [38]. Chimeras, mitochondrial, and chloroplast sequences were removed. Alpha diversity analyses of fungal and bacterial communities were performed using phyloseq [39] and vegan [40].

## Results

### The biofertilizer exhibited distinct diversity patterns between fungi and bacteria

We processed amplicons of the ITS region and 16S rRNA gene V3-V4 region from substrate and plant roots samples derived from the biofertilizer. Our analysis revealed 62,587 fungal ITS sequences associated with the substrate and 56,566 associated with plant roots. After filtering out singletons, we identified 635 fungal OTUs in the substrate and 576 in the plant roots. It indicated that the fungal diversity in the substrate (H’=4.5) was higher than that in the roots-associated fungi (H’=3.4; Table 1). In addition to fungal diversity, we found several OTUs from plants and microeukaryotes, suggesting a more complex community (S1 Fig).

**Table 1.** Sequencing outputs and alpha diversity indexes.

|  | Substrate-associated fungi (ITS) | Roots-associated fungi (ITS) | Substrate-associated bacteria (16S rRNA gene) | Roots-associated bacteria (16S rRNA gene) |
| --- | --- | --- | --- | --- |
| Raw paired sequences | 62,587 | 56,566 | 842,909 | 902,266 |
| OTUs (97%) | 617 | 374 | 6,963 | 8,808 |
| Chao 1 (±SE) | 625 ± 4 | 429 ± 14 | 7,350 ± 29 | 8,971 ± 16 |
| Shannon ($H'$) | 4.6 | 2.5 | 6.08 | 7.62 |
| Simpson | 0.97 | 0.83 | 0.98 | 0.99 |

Regarding the bacterial 16S rRNA gene amplicons, we got 842,909 sequences from the substrate and 902,266 sequences from the roots. After processing sequences, we found 6,971 OTUs in the substrate and 8,817 OTUs associated with the root samples. Regardless of sample type, bacterial communities were more diverse (H’=6.08-7.62) than fungal communities (H’=3.4-4.5; Table 1). Opposite to fungal communities, diversity from the roots (H’=7.62) was higher than from the substrate (H’=6.08). Together, these results indicate that using substrate and roots maximizes biofertilizers’ fungal and bacterial diversity.

### Microbial compositions of biofertilizer communities

The predominant phylum from fungal communities was Ascomycota, followed by Glomeromycota, Basidiomycota, and Mucormycota (Fig 2). However, their relative abundance (RA) changed depending on the analyzed sample. Although Ascomycota is in a high proportion (RA=0.54) in the substrate samples, the phylum was more abundant in the roots (RA=0.86). Glomeromycota and Basidiomycota were more abundant in the substrate (RA=0.23 and 0.16) than in the roots, where they were drastically reduced (RA=0.01 and 0.03). Conversely, Mucoromycota was less abundant (RA=0.05) in the substrate than in the roots (RA=0.09).

**Fig 2.**
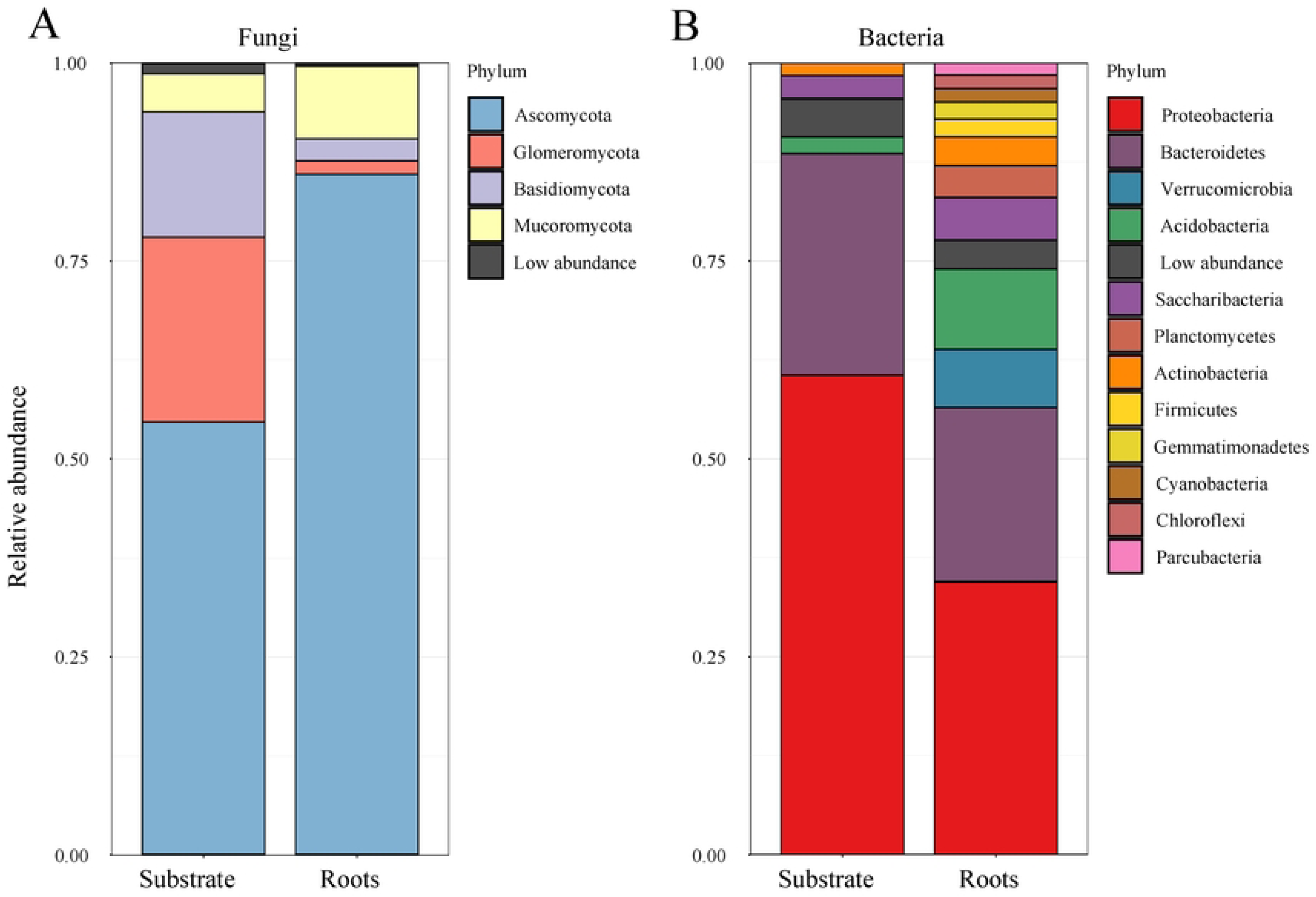
Phyla diversity of fungal (A) and bacterial communities (B) from the substrate or plant roots-associated.

Bacterial communities were dominated by *Proteobacteria*, *Bacteroidetes*, *Verrucomicrobia*, *Acidobacteria*, *Saccharibacteria*, and *Actinobacteria* (Fig 2B). In the substrate, the highly abundant phyla *Proteobacteria* (RA=0.605) and *Bacteroidetes* (RA=0.279) dominated, whereas *Saccharibacteria* (RA=0.029), *Acidobacteria* (RA=0.021), and *Actinobacteria* (RA=0.016) were in low abundance. In the roots, *Proteobacteria* (RA=0.344) and *Bacteroidetes* (RA=0.219) reduced compared to the substrate, while (RA=0.036) increased. *Verrucomicrobia* was negligible (RA=0.013) in the substrate but in a considerable abundance (RA=0.073) in the substrate (Fig 2).

The ITS region sequence analysis allowed us to identify 182 fungal species. Of them, 98 were exclusive from the substrate, 11 from the plant roots, and 73 were shared between both samples (Fig 3A, upper section). *Bipolaris sp.*, *Rhizopus arrhizus, Scutellospora heterogama*, *Pyrenochaetopsis leptospora*, *Aspergillus subversicolor*, *Pseudotomentella alobata*, *Talaromyces marneffei*, *Edenia gomezpompae*, *Mycosphaerella tassiana*, *Diversispora celata*, *Didymella exigua*, *Curvularia lunata*, *Rhizopus microsporus*, *Atractiella rhizophila, Bipolaris sorokiniana*, and *Gigaspora margarita* were the more abundant fungal species (RA > 0.01) in the substrate and the roots (Fig 3; S1 Table). The 16S rRNA gene analysis revealed 964 bacterial genera, of which 80 were exclusive from the substrate, 269 were exclusive from the root, and 615 were found in both samples (Fig 3B, upper section). *Parabulkholderia-Burkholderia*, *Rhizobium*, *Sphingomonas*, *Chitinophada*, *Mucilaginibacter*, *Bradyrhizobium*, *Taibaiella*, *Flavosolibacter*, *Opitutus*, *Acidibacter*, *Bacillus, Haliangium*, Flavitelea*, Bryobacter*, *Niastella*, *Filimonas*, *Massilia*, *Dyella*, as well as several uncultured bacteria, were dominant in the substrate or the roots (RA > 0.01; Fig 3; S2 Table). The complete lists of taxa from fungal and bacterial communities are available in supplementary tables (S1 Table and S2 Table).

**Fig 3.**
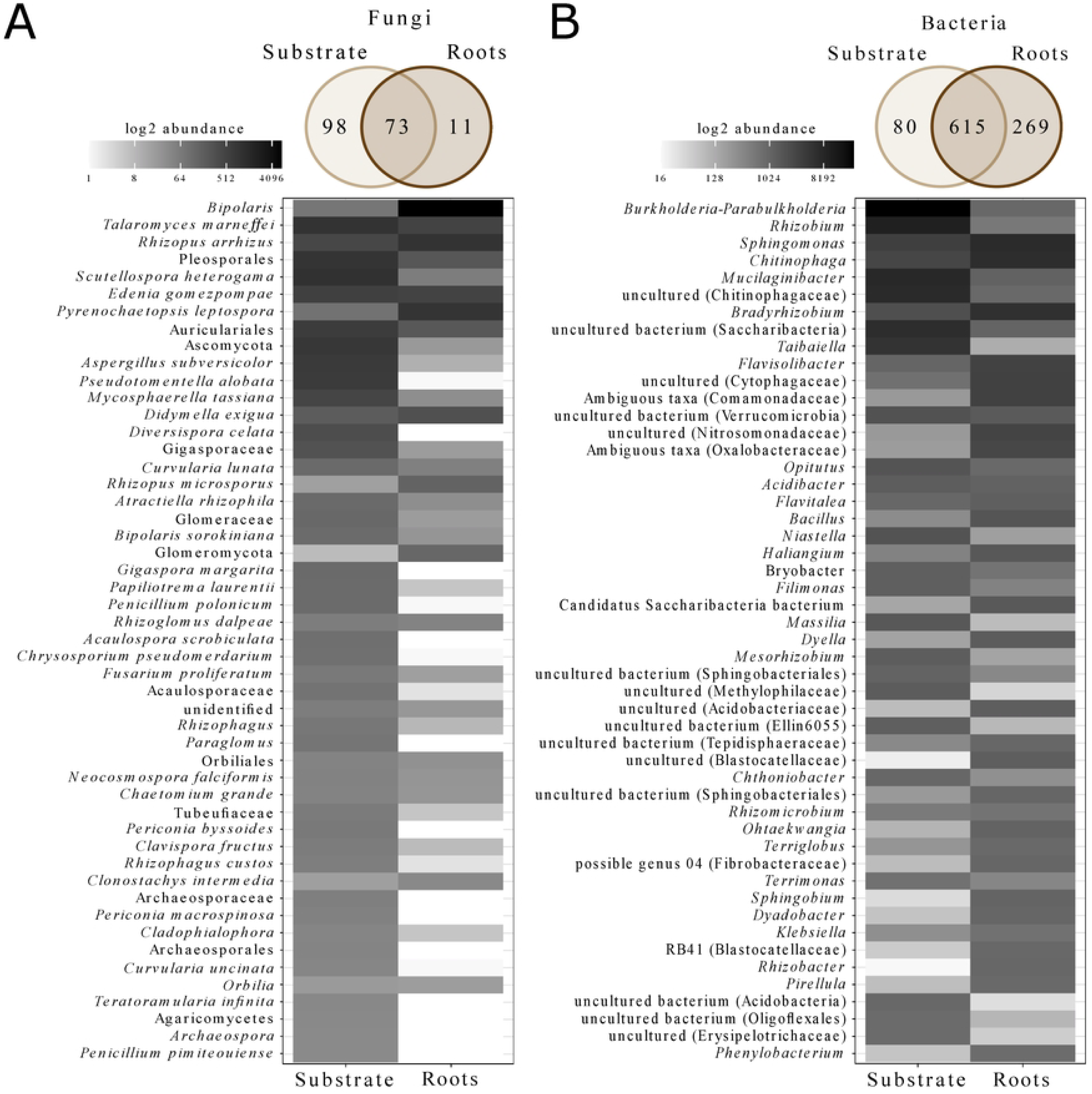
Microbial consortia in biofertilizers. The fungal species (A) and bacterial genera communities (B) present in the biofertilizer are depicted in Venn diagrams, illustrating the number of taxa exclusive to, and shared between, the substrate and the plant roots. Additionally, heat maps display the relative abundance of the most prevalent taxa.

## Discussion

### Assembly of microbial communities

This work evaluated the diversity levels of the biofertilizer microbial communities, showing that its production optimizes the fungal and bacterial communities containing plant growth-promoting microorganisms. While bacterial diversity was increased between the substrate and the roots, fungal diversity followed an inverse pattern (Table 1). The increase in diversity in root bacterial communities (Table 1) may be attributed to the nutrient-rich environment established in the rhizosphere by the plant metabolite secretion [41]. Previous studies have shown that rhizospheres host higher bacterial alpha diversity than soils [42,43].

A common garden experiment found that the microbiomes of ruderal plants had higher alpha diversity than their soils. Similarly, when the same soils were tested on domesticated plants, the alpha diversity decreased in the rhizosphere, suggesting that the rhizosphere provides a microenvironment that supports bacterial diversity [42]. Our findings suggest that the sandy substrate will serve as a source of bacterial inoculum that is enriched in diversity by the presence of plant roots, which act as a system for attracting and promoting bacterial growth. Notably, specific genera such as *Sphingomonas*, *Flavisolibacter*, and *Opitutus* were more abundant in the roots than in the substrate. Additionally, some *Rhizobiales* were exclusively found in the roots (Fig 3B; S2 Table). Since *C. juncea* and *C. ensiformis* are legumes, the root-exudated flavonoids could attract exclusive rhizobia from the roots to induce nodulation [44]. Additionally, some bacteria are vertically inherited, such as *Sphingomonas* found in several generations of *Crotalaria pumila* seed microbiome [45].

The reduction of the root fungal diversity may be explained by the selection process driven by the roots. In which inoculated fungi are selected by their affinity with root-released metabolites and host genotype [41,46]. We recognized 182 fungal species in the inoculated substrate. However, only 84 were detected in the roots (Fig 3A), similar to the previously observed reduction of the fungal diversity between the soil and the root found by other works [47,48]. Although plants like *Brachiaria*, *Crotalaria,* and *Canavalia* are known for establishing arbuscular mycorrhizal interactions that improve their growth under unfavorable conditions [49–52], we did not detect *D*. *celata* and *G*. *margarita* in the roots (Fig 3A; S1 Table; Table 2). The low abundance of Glomeromycota associated with roots is a common pattern found in several cultivated species, such as agaves [53], sugarcane [54], cactus [55], and wheat [49]. Nonetheless, the substrate kept a high proportion of arbuscular fungi from this phylum compared with soils [56,57], reflecting a thriving selection of fungal symbionts other plant species can recruit.

**Table 2.**
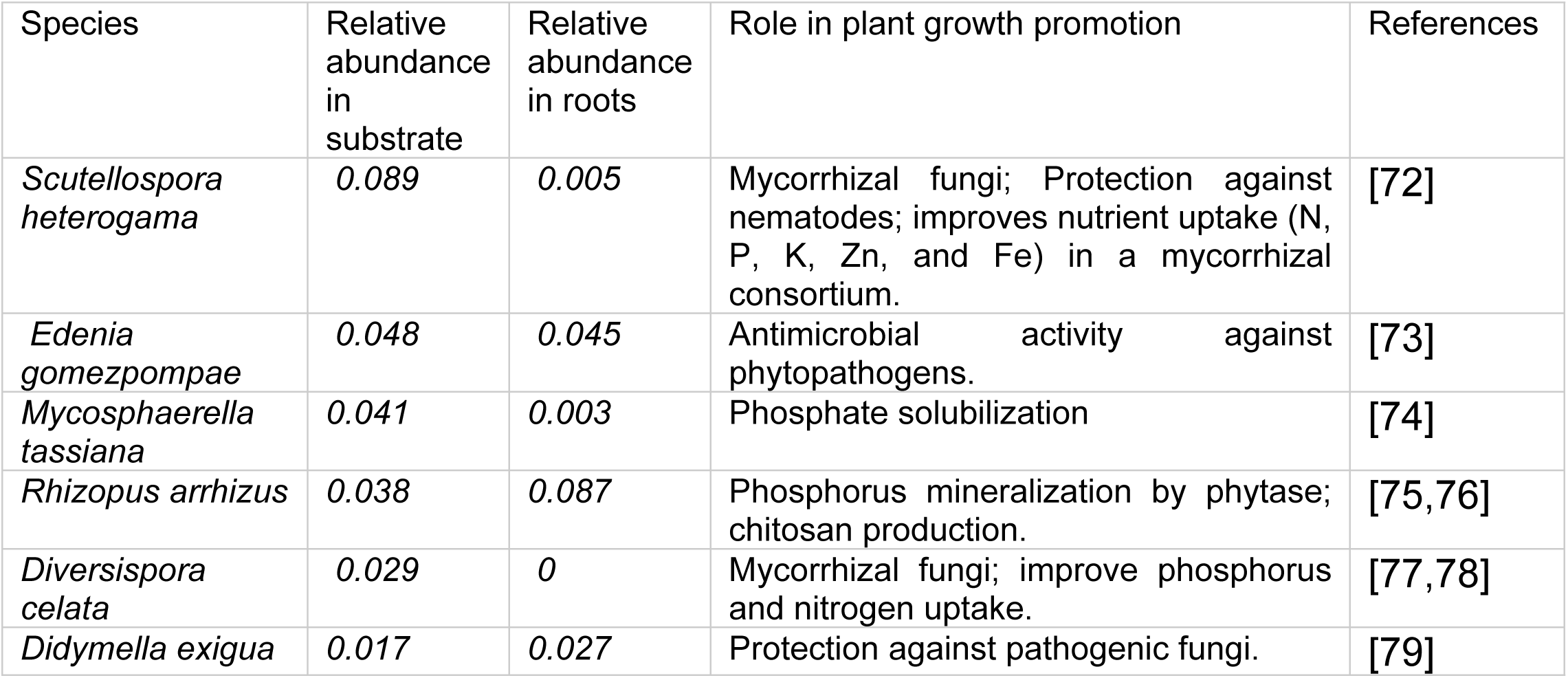
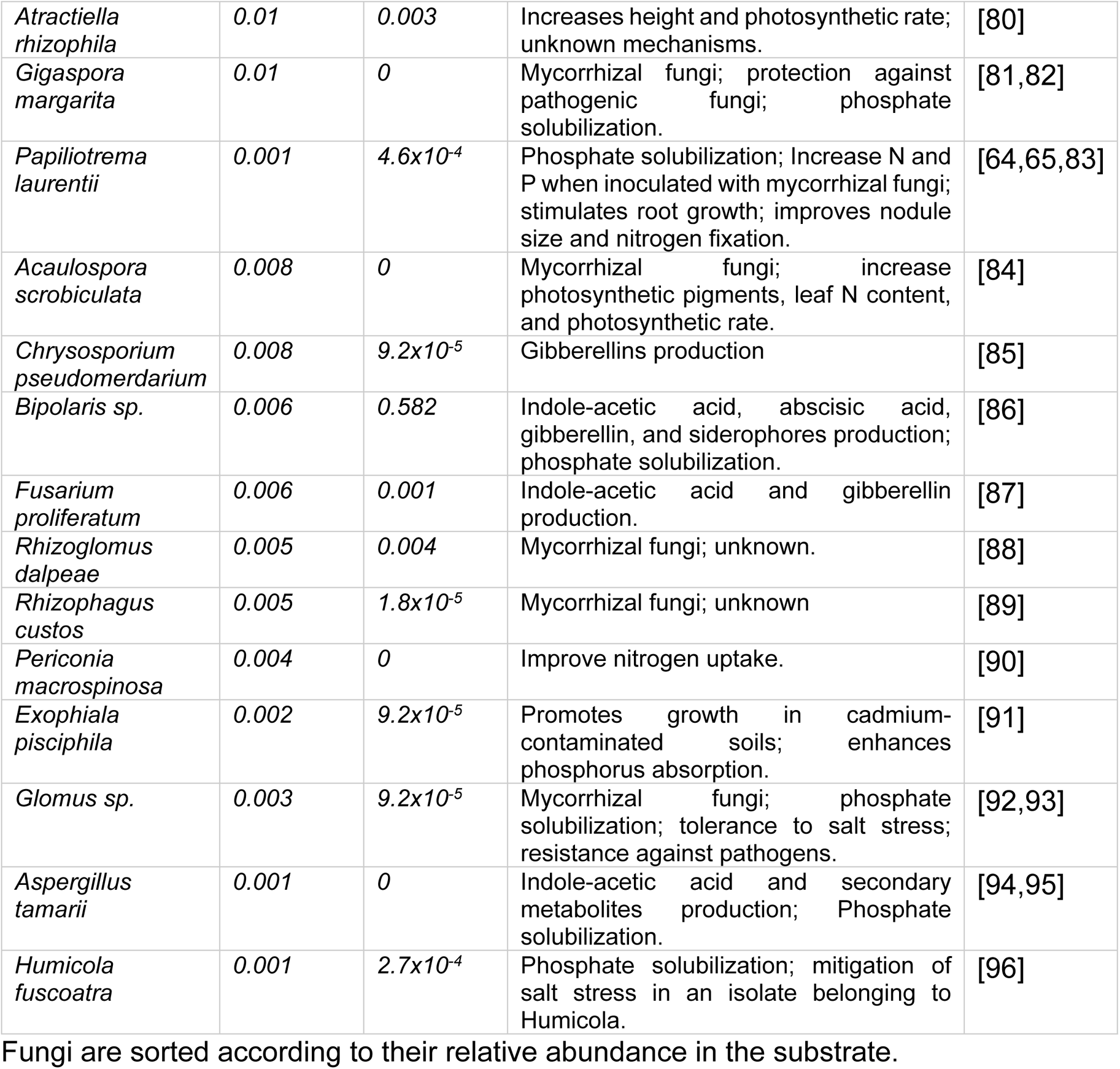
Fungi with a possible role in plant growth promotion in the biofertilizer

It is essential to highlight that we divided the samples of substrate and roots to understand plant-microbe interactions in line with other works [48,58] as a process of microbial colonization from soil to roots [46]. Despite the bioinoculant being composed of dried roots, particles of the substrate can adhere to them during processing and play a role in the observed positive effects on plant phenotype. Therefore, a microbial characterization of the biofertilizer should consider the beneficial microbes in both the substrate and the roots.

We present a model that explains the assembly of bacterial and fungal communities in the biofertilizer (Fig 4). The inoculation of mycorrhizal fungi adds bacteria to the sandy substrate. During the plant growth process in greenhouses, beneficial microorganisms are intentionally added to biofertilizers, and some microbes are inadvertently introduced through natural means. For instance, other unintentional microbial inoculations can occur through watering, which often uses non-sterile water, or environmental aerosols and dust. When the plants are introduced to the substrate, they filter the fungi based on their ability to interact with *B. brizantha, C. juncea*, and *C. ensiformis*, reducing fungal diversity in the root-associated communities. However, plants exude metabolites to the substrate, creating a nutrient-rich niche that boosts bacterial diversity [41,59]. Although the root-associated communities contain bacteria and fungi, our results suggest that incorporating the substrate fraction to the biofertilizer is relevant to maintain the whole diversity of the original fungal community. Fungal diversity from the substrate includes arbuscular endomycorrhizal fungi with agricultural relevance from the phylum Glomeromycota [60]. For this reason, both the substrate and the roots are essential to the biofertilizer since their synergistic effect provides a broad spectrum of microbes with the capacity to promote plant growth in several plant species. In addition, the biofertilizer process resembles host-mediated microbiome engineering (HMME). HMME is a multigenerational process that focuses on the host and sub-selects beneficial microbes at a community level rather than individually. Since the HMME can be targeted to increase plant tolerance to environmental stress [61], new biofertilizers could be designed by trap cultures and the positive selection of stress-resistant phenotypes.

**Fig 4.**
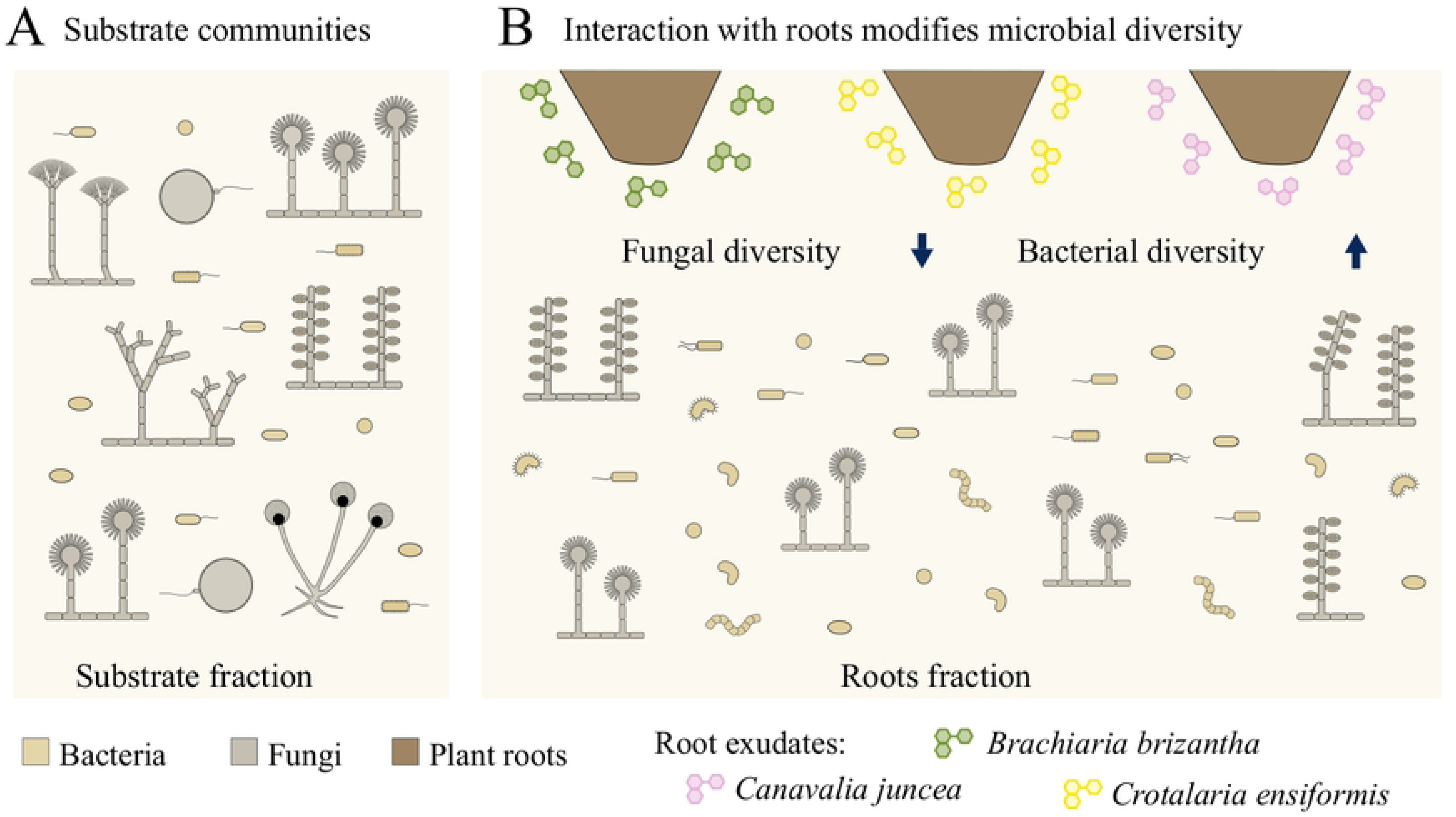
A comprehensive model explains the biofertilizer’s observed bacterial and fungal diversity. (A) Initially developed for fungi and bacteria, the consortia formation process involves introducing bacteria to the substrate through the initial mycorrhizal fungi inoculum. (B) Subsequently, the plants actively select their fungal partners, reducing fungal diversity near the roots. Interestingly, the root-associated bacteria displayed greater diversity, likely due to the nutrient-rich environment created by the plant root exudates.

### Microbes with reported plant growth promotion activity

According to the literature, ITS and 16S rRNA gene sequencing allowed us to identify taxa that may be responsible for plant growth promotion (Table 2; Table 3). Arbuscular mycorrhizal species such as *Scutellospora heterogama*, *Diversispora celata*, and *G. margarita* were predominantly found in the substrate (Fig 3; Table 2; S1 Table). Besides the direct benefits to plants, mycorrhizal fungi probably contribute to establishing other plant growth-promoting microorganisms [62,63], found in higher abundance in (Fig 3; Table 2; Table 3; S1 Table). *Papiliotrema laurentii* can interact with *Glomus mossae* to enhance nutrient content in roots and leaves [64] or with other mycorrhizal species to increase nodule size and nitrogen fixation [65].

**Table 3.** Bacteria with a possible role in plant growth promotion in the biofertilizer

| Genus | Relative abundance in substrate | Relative abundance in roots | Role in plant growth promotion | References |
| --- | --- | --- | --- | --- |
| *Sphingomonas* | 0.029 | 0.058 | Indole acetic acid and gibberellin production. | [97] |
| *Bacillus* | 0.008 | 0.010 | Nitrogen fixation, phosphate solubilization, induction of iron acquisition genes from plants; alteration of plant growth hormone homeostasis; drought and salt stress resistance; production of antimicrobial compounds; induced systemic resistance. | [98] |
| *Chitinophaga* | 0.006 | 0.009 | Phosphate solubilization; production of chitinase against pathogenic fungi. | [99,100] |
| *Bradyrhizobium* | 0.055 | 0.009 | Nitrogen fixation; indole acetic acid production; phosphate solubilization. | [101,102] |
| *Burkholderia* | 0.225 | 0.007 | Nitrogen fixation, phosphate solubilization; indole acetic acid production; antifungal activity. | [103] |
| *Klebsiella* | 6.9x10<sup>-4</sup> | 0.008 | Indole acetic acid and siderophores production. | [104] |
| *Rhizobium* | 0.087 | 0.004 | Nitrogen fixation, phosphate solubilization, indole acetic acid, gibberellin production, and induced systemic resistance. | [101,105,106] |
| *Mesorhizobium* | 0.009 | 0.003 | Nitrogen fixation; phosphate and potassium solubilization; Indole acetic acid and siderophores production; | [107] |
| *Massilia* | 0.011 | 0.0013 | Indole acetic acid and siderophore production. | [108,109] |
| *Sphingobium* | 0.007 | 8.2x10<sup>-4</sup> | Improve tolerance to cadmium | [110] |
| *Dyadobacter* | 0.007 | 4.3x10<sup>-4</sup> | Nitrogen fixation; phosphate solubilization | [111,112] |
| *Dyella* | 0.011 | 3.5x10<sup>-4</sup> | Phosphate solubilization | [113] |

Bacteria are sorted according to their relative abundance in the roots.

Other highly abundant fungal species are *Talaromyces marneffei* and *Aspergillus subversicolor* (Fig 3 and S1 Table). To our knowledge, *T. marneffei* is a human pathogen [66] with no records in the plant microbiome. However, some *Talaromyces* species enhance plant growth by controlling pathogens [67,68][59,60] or producing antioxidant enzymes and osmolytes [69]. Although *A. subversicolor* was isolated from coffee [70], there is little information about the species. On the other hand, some species of *Aspergillus* can benefit agricultural production due to their ability to solubilize and mineralize phosphorus and produce secondary metabolites and phytohormones [71].

We suggest that biofertilizer improves plant growth through four main mechanisms: nutrient uptake, phytohormone production, stress tolerance, and resistance to pathogens (Table 2; Table 3). The primary mechanism for plant growth promotion by fungus seemed related to phosphorus acquisition mediated by phosphate solubilization [82] and phosphate mineralization [75]. Although we identified some phosphate-solubilizing bacteria [106], our dataset suggests that while fungi are mainly involved in phosphorus nutrition, bacteria may play a key role in nitrogen uptake by nitrogen fixation [101,107,114]. Both fungi [115] and bacteria [107,108] can produce siderophores for iron uptake. Regarding stress response, there are reports of microorganisms, such as *Glomus* and *Bacillus,* involved in salt and drought stress tolerance [93,98]. *Bacillus subtilis* increases the tolerance of plants to salt and drought stress by modulation of abscisic acid, one of the main phytohormones for stress response [98]. We found several microorganisms involved in resistance against pathogens. For instance, *E. gomezpompae* produces naphthoquinone spirochetal, secondary metabolites with antifungal activity against the plant pathogens *Phytophthora capsici, P. parasitica,* and *Fusarium oxysporum* [73].

Finally, it is necessary to mention that some taxa we considered possible plant growth promoters have species with pathogenic activity. For example, some *Bipolaris* and *Fusarium* species are responsible for diseases that can cause rot in plant organs [116,117]. However, we found that some species of *Bipolaris sp. CSL-1* produces indole acetic acid and gibberellins, increasing seedling biomass and chlorophyll content [86]. Some *Fusarium* species could promote growth by phosphate solubilization, synthesis of phytohormones, and siderophore production [96,118]. Alternatively, several biofertilizer microbial communities could produce metabolites to antagonize pathogens from the same community [93,98,119]. Previous works suggest that antagonistic interactions between bacteria and fungi may promote plant growth [120], and resource competition between closely related species (non-pathogenic vs. pathogenic) may also exclude pathogens from plant roots [121]. In addition, PGPB consortia are composed of mutualistic organisms and contain microbes without directly benefiting plants that play essential roles in their communities [122].

## Conclusions

We investigated the fungal and bacterial diversity of a commercial biofertilizer, which consists of plant roots added to a mycorrhizal-inoculated substrate. Our study identified 182 fungal species and 964 bacterial genera. The dominant fungi were *Bipolaris* sp, *Rhizopus arrhizus*, and *Scutellospora heterogama*, while the dominant bacteria were *Burkholderia*, *Rhizobium*, *Sphingomonas*, and *Chitinophaga*. Interestingly, these microbes are known to promote plant growth. We also found that fungal diversity was higher in the substrate, while bacterial communities were more diverse in the roots. Our results suggest that initial inoculation provides a high fungal diversity, while plant incorporation diversifies bacterial communities, creating a range of microorganisms that promote plant growth. Moreover, the long-term selection of beneficial microbial communities that interact with plant roots and improve their phenotype can create new biofertilizers that address specific issues such as biotic and abiotic stress.

## Acknowledgments

This work was supported by Universidad Nacional Autónoma de México by the projects DGAPA-PAPIIT-UNAM IN221420 to LDA. Conacyt Ph.D. scholarship (CVU 742786) to CHA.

## Supporting Information

S1 Fig. Eukaryotic diversity. A) Phyla diversity and B) Relative abundance of the non-fungal genus.

S1 Table. List of fungal species. Species are sorted according to their relative abundance in the substrate.

S2 Table. List of bacterial genera. Genera are sorted according to their relative abundance in the roots.

## Notes

### Competing Interest Statement

I have read the journal's policy and the authors of this manuscript have the following competing interests: LDA is an Academic Editor for this journal. This does not alter our adherence to PLOS ONE policies on sharing data and materials. DTA is the designer of the commercially available biofertilizer used in this study. DTA provided biological material and valuable input during manuscript development. However, she did not have a direct role in the study design, data collection, analysis, or interpretation of the results.

